# Quantifying free behavior in an open field using k-motif approach

**DOI:** 10.1101/735399

**Authors:** Marein Könings, Mark Blokpoel, Katarzyna Kapusta, Tom Claassen, Jan K. Buitelaar, Jeffrey C. Glennon, Natalia Z. Bielczyk

## Abstract

Quantification and parametrization of movement in animal models is widely used in behavioral paradigms. In particular, free movement of an animal in controlled conditions (e.g., the open field paradigm) is used as a proxy for indices of baseline and drug-induced behavioural changes. However, the analysis of this is often time- and labour-intensive and existing algorithms do not always classify the behaviour correctly.

Here, we propose a new approach to quantify behaviour in an unconstrained environment: searching for frequent patterns (k-motifs) in the time series representing position of the subject over time. Validation of this method was performed using subchronic quinpirole-induced changes in open field experiment behaviors in rodents. Analysis of this data was performed using k-motifs as features to better classify subjects into experimental groups on the basis of behavior in the open field. Our classifier using k-motifs gives as high as 94% accuracy in classifying repetitive behaviour versus controls which is a substantial improvement compared to currently available methods including using standard feature definitions (depending on the choice of feature set and classification strategy, accuracy up to 88%). Furthermore, vizualization of the movement / time patterns is highly predictive of these behaviours. By using machine learning to create features in a data driven fashion, this can be applied to general behavioural analysis across experimental paradigms beyond the open field.

## 1. Introduction

### 1.1. Open field paradigms in translational psychiatry

Rodent behavioural research analyses patterns of behaviour as face and predictive validity measures for human psychiatric and neurological conditions.These behavioural paradigms are relevant to a number of cognitive disorders, including schizophrenia, depression, bipolar disorder[1], anxiety and autism[2]. The open field paradigm is commonly deployed for the study of movement, learning, sedation and anxiety related aspects[3]. In the open field paradigm, an animal is placed in a constrained environment (a well-lit circular, square or rectangular area, bounded either by insurmountable walls or deep gaps) which it is free to explore. Its behavior is then recorded over time. The characteristics of the environment can differ between experiments and may deploy objects to study interaction between the animal and object to examine reactions to novel and stressful situations. An open field with objects can also be used to examine repetitive checking behaviour, an important aspect of obsessive compulsive behaviors[4]. Commonly recorded variables include horizontal locomotion (based on a count of transitions between marked areas within the field, Figure 2), vertical activity (based on the animal’s rearing and leaning behaviour), as well as the latency (time) spent in certain areas and more specific behaviours such as head shakes and grooming[3]. Automated systems record all of the necessary variables with high precision and flexibility[5], by recording the animal’s position using a video camera and processing the resulting footage with specialised software such as Noldus EthoVision 3.0[6]. The open field is also often divided into a grid of zones for the analysis which can be of differing sizes dependent on the size of the open field. Typically this is divided into 5 by 5 zones. A range of experimental designs are used within the open field paradigm. Subjects are divided into experimental groups which undergo different interventions. Differences between groups in terms of observed open field behaviour is then assumed to result from the difference in intervention deployed. These experimental interventions include changes to the objects in the open field, introduction of another animal, differences in the amount / type of food available or administration of drugs expected to induce behavioural changes[7].

### 1.2. Automated methods for quantifying rodent behaviour

In order to accurately assess the differences in behaviour between experimental groups, the objective quantification of the behaviour is important. Although currently available software such as Noldus Theme is able to extract raw behavioural variables from video footage, little has been done to automate and optimize further analysis of the collected datasets. These approaches detect *T-patterns*, which denote the repeating sequences of fields visited by the animal during the experiment[8, 9]. This method while useful also involves a large number of significance tests, thereby increasing the risk of false positive results[10]. In addition, the T-patterns method assumes that all areas of the open field are equally likely to be visited. In reality, the opposite is true: some areas are visited far more often than others. Furthermore, zone transitions are computed using the T-pattern method with no regard to the temporal relation between crossings. That is, the time that a subject spends within a zone is of no consequence, only the order in which zones are visited counts. This is problematic as this does not enable a distinction between behaviours that are time-sensitive (e.g. spending time near an object as opposed to just passing by it). This is a consequence of the T-pattern method performing only spatial and not temporal segmentation. Finally, the T-pattern method only utilises the absolute position of the subject, and does not account for other factors such as objects in the open field or the distance between the animal and the open field boundary. As such, behaviours dependent on the relative location of these objects cannot be detected using T-patterns. If the location of an object changes, for example between subjects, the T-pattern method cannot discover a pattern if both subjects display the same behaviour relative to the object but in different absolute positions. Lastly, the T-pattern method is only available as a part of a commercial software, *Theme* by *Pattern Vision* (http://patternvision.com/products/theme/), which is a black-box software with a basic interface, and does not allow for accessing the source code.

### 1.3. The k-motif approach

Here we sought to design a new method for quantifying behaviour in the open environment. In an attempt to overcome the above issues, we proposed a method which takes into account the temporal dynamics of the movement as well. This method adapts the k-motifs algorithm to search for the hierarchical structure of repeating patterns in behaviour. The most common patterns were then used to train a classifier to distinguish between experimental groups between quinpirole and vehicle treated rats in the open field. Here, we improve the classification of experimental groups (further referred to as classes) on the basis of behavioural differences in the open field.

In section 2.1, we provide a detailed description of the experimental datasets used for developing the classifier. In section 2.2, we introduce the k-motifs approach. Then, in section 2.5, we report the results of classification with use of k-motifs as features, and compare this classification performance with classifiers using standard features. Lastly, in section 3, we critically discuss the results and potential applications for this methodology. We made all the tools developed for this paper accessible in open source.

We dedicate this work to researchers looking for data-driven biomarkers of behavior in the open field. These patterns can help in better understanding of the characteristics of disorders related to movement, such as Obsessive-Compulsive Disorder, Tourette Syndrome, Parkinson’s Disease or Hutchinson’s Disease. The k-motif method proposed in this manuscript can also help to measure effects of pharmacological therapy with higher sensitivity as the performance in classification between experimental and control group achieved with our method is the highest among all the available methods.

## 2. Materials and methods

### 2.1. The experimental datasets

The experimental datasets includes open field recordings in rats, collected at the Radboud University Nijmegen Medical Centre under ethical approval number DEC-2012-281. The sample included 6 subjects in the experimental group and 6 controls. All subjects were male Sprague Dawley rats (Charles River, Germany). The open field setup includes an open square area of size 160 × 160 [cm], placed 60 [cm] from the ground. An additional virtual circuit 20 [cm] wide around the table is added to the open field size in order to record behaviour where the subjects’ head or tail reaches out over the table. The resulting area of 200 × 200 [cm] is divided into 25 40 × 40 [cm] zones. The area contained four small fixed objects (cubes or cylinders) to encourage exploration in animals (Figure 1 A, B). The objects were distributed as in[11], and included two black and two white objects. The experimental sample included 6 subjects in an experimental intervention group and 6 controls. In total, data was recorded from subjects across 13 30-minute sessions in the open field, which amounts to a total of 156 recorded sessions. Injection and training sessions took place every day for 13 consecutive days. Prior to each open field session, the subjects within the experimental intervention group received an injection of quinpirole (0.5mg/kg i.p.). The behaviour in the open field was recorded with an overhead video camera. The resulting footage was then processed by a video tracking system Ethovision 3.0 from Noldus Information Technology BV, the Netherlands (Wageningen, The Netherlands) [6], which produced multivariate time series for each 30-minute session, representing multiple variables (Table 6, Supplementary Material 4.1). The set of available variables is represented here by *V*. The collection of time series resulting from processing a video footage from one session in one subject, corresponds to a single *d* in *D* (Equation 1). Examples of the visualisations of the movement are presented in Figure 1. The datasets were preprocessed according to a preprocessing scheme is introduced in Supplementary Material 4.2).

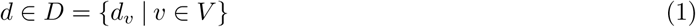

**Figure 1:**
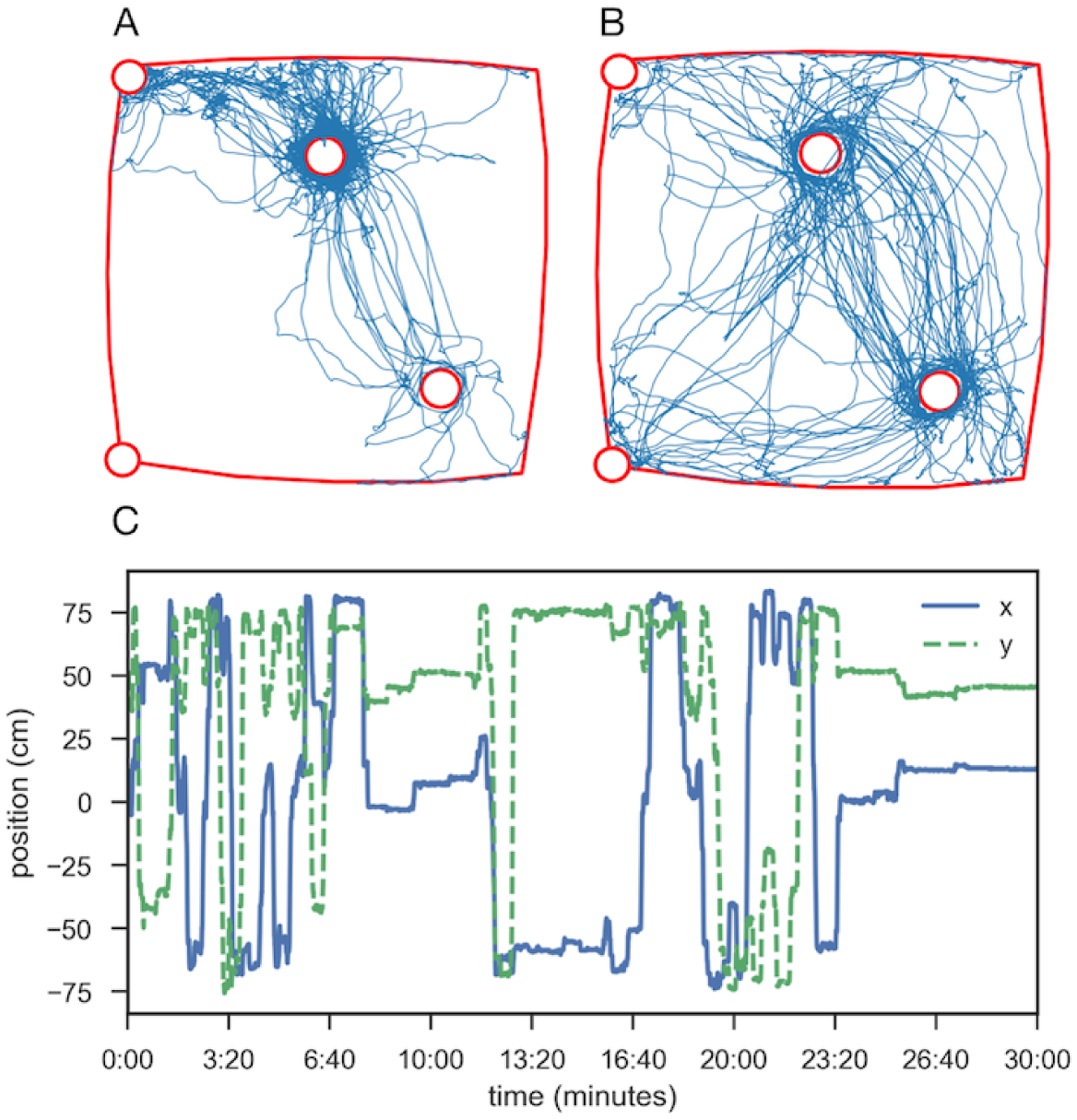
Movement of example subjects during single sessions. Subfigure **A** shows the spatial, ‘top-down’ view of the path taken by an exemplary subject. The borders of the open field are shown in red, with the four objects shown as red circles. The same path’s *x* and *y* coordinates over time are shown in temporal subfigure **C**. Subfigure **B** shows the path taken by an exemplary subject.

### 2.2. K-motif method

In an effort to address the limitations of T-patterns mentioned in Section 1, in this work, the *k*-motifs method is employed for feature extraction: finding recurring patterns [12, 13] in the data. This method involves subsequent use of two algorithms: Symbolic Aggregate Approximation (SAX, [14]) and Sequitur grammar induction [15]. The first part, SAX, segments the data both in the temporal domain and in the spatial domain. The second part, grammar induction using Sequitur, finds patterns in the resulting approximation by constructing a hierarchical grammar. The *k*-motifs algorithm successfully yields frequent patterns from a time series, which can be used to characterise the data. However, to enable classification of time series data, each series must be reduced to a feature vector. A feature vector is a list of numbers of a set length, where each number describes an aspect of the data. These numbers can then be compared between series, to determine their relative similarity. In order to use the patterns extracted by *k*-motifs, a conversion strategy to feature vectors must be devised. Additionally, while patterns in raw movement data are useful, even more patterns may be found by exploiting the animal’s position relative to different aspects of the open field. Finally, *k*-motifs produces a very large number of patterns, so in order to ensure the conciseness of the quantification method, some subset of the patterns must be selected for the final feature vector. All these topics will be discussed in the following sections.

#### 2.2.1. Symbolic Aggregate Approximation (SAX)

The Symbolic Aggregate Approximation (SAX) algorithm [14] takes a real-valued time series *x*(*t*) and *discretises* the time series, starting by slicing the values into segments of width *w*. For each segment, the values contained within are averaged. This leads to a new time series *t*(*t*) of length 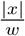. Note that if the length of *x*(*t*) is not divisible by *w* (length of *x*(*t*) is not a multiple of *w*), *x* is padded with the last value of *x* until its length is divisible by *w*.

Next, the *range of the values* of *t*(*t*) is divided into a number of sections (further referred to as ‘bins’). The bins are defined by means of z-scores from a normal distribution with mean and standard deviation derived from the original data *x*(*t*). We denote the number of bins by *α*, and each bin is represented by the lower limit in its value range. For example, if the values fall between −4.0 and 4.0, and *α* = 4, the bins are: [−4.0, −2.0] represented by −4.0, [−2.0, 0.0] represented by −2.0, [0.0, 2.0] represented by 0.0, and [2.0, 4.0] represented by 2.0.

Additionally, the bins are given unique labels. In this work, bins are labeled from 0 to *α* − 1, using the ordering of the bins from the lowest to the highest. Then, each value in *t*(*t*) is replaced by the label of the associated bin to which is belongs - namely, the highest bin whose lowest value is lower than the given value in *t*(*t*). The obtained result is an approximate symbolic representation of the original time series. This transformation is formalised in Equation 2. An example segmentation is demonstrated in Figure 2 A.

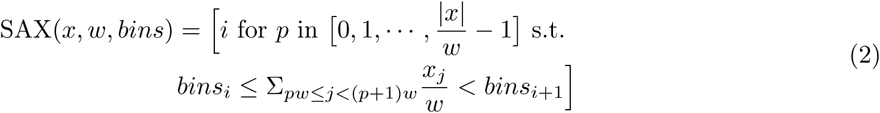

**Figure 2:**
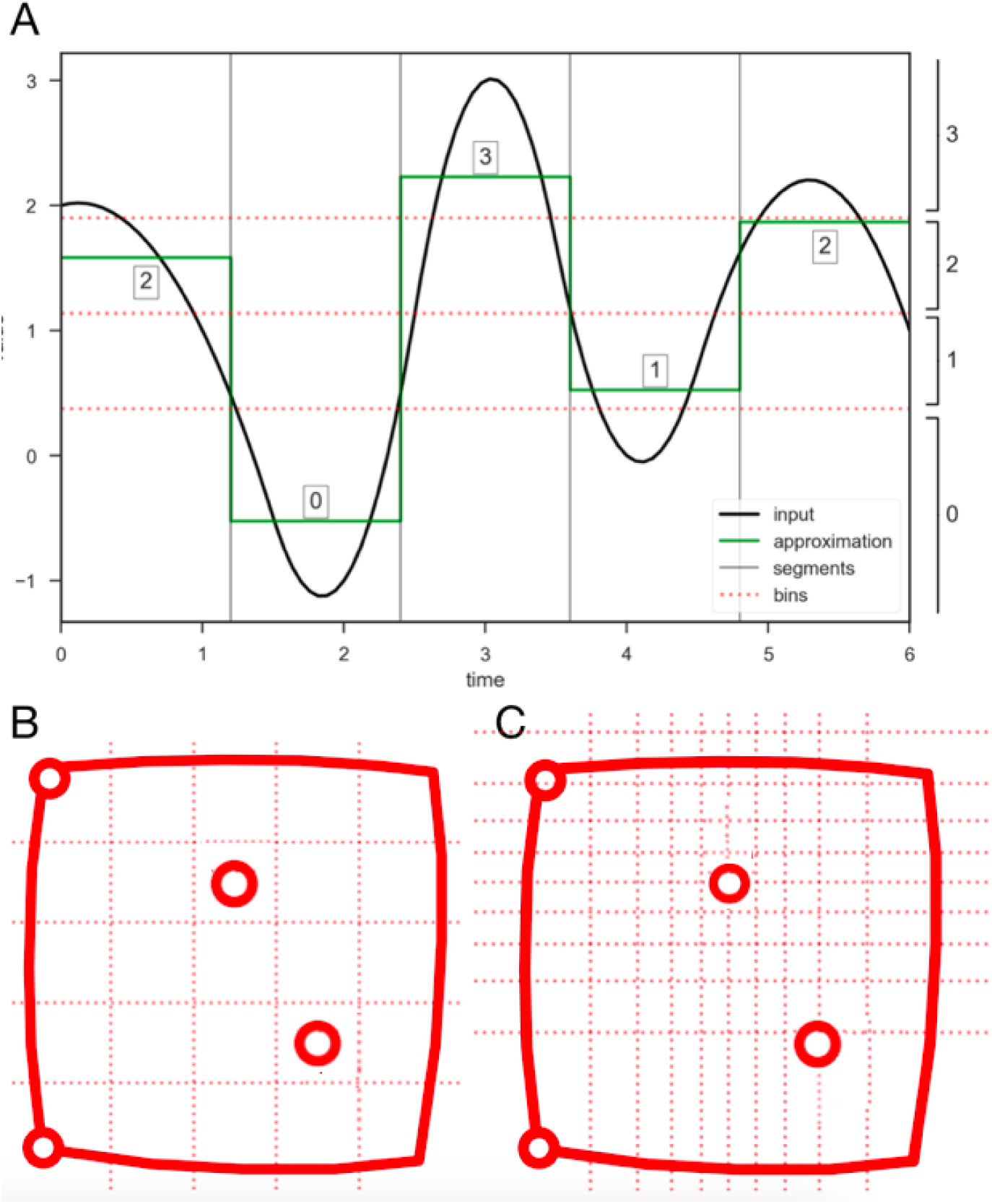
The SAX algorithm [14]. **A**: the real-valued input data is presented in black. It the time resolution of the recording is high, this may be a near-continuous sequence. In this example, parameter *w* is set to such a value that the input is divided into segments of 1.2 time units. The mean value of the time series within each segment is used to produce the approximation sequence (presented in green). Finally, for each mean value, the corresponding bin is determined. In this example, there are four bins (*α* = 4) delineated using dotted red lines. Bins width is determined in terms of standard deviations from the mean value. The final output in this example, is a sequence of labels 2 0 3 1 2. **B**: standard division into 25 zones (as implemented in the T-pattern method). **C**: the division of open field with use of SAX (*α* = 10, which means 10 bins along each dimension), with all positional data. Note the underlying normal distribution of visits is reflected in the sizing of the bins. In less popular areas, larger bins are available and less detail is recorded.

Assuming normal distribution of the input datasets, this ensures that each value in *t* is equally likely to fall into each bin, and thus that each event (falling into a bin 0 to *α* − 1) occurs equally often. This is explained in Figure 2, where *α* = 4. In this example, the value range in the time series, is dissected into four equiprobable bins, centred on the mean of the time series. Note that, for any considered variable (e.g., the animal’s *x* position on the open field table), the bins are precomputed using *all* available datasets: the bins are defined for the merged datasets from all the subjects within the experimental group. The purpose of this step is to have one, uniform output sentence for all subjects. An example set of bins, computed for all two-dimensional positional data in our datasets, is presented in Figure 2 B. An advantage of using bins based on the normal distribution derived from the datasets - rather than equally-sized bins - is that the proportion of visits in all bins becomes more uniform^1^.

In order to apply the SAX algorithm, we reimplemented the original version of the algorithm using Python, the code is included in the open GitHub repository, at https://github.com/MareinK/kmotifs-paper-code.

#### 2.2.2. Grammar induction using Sequitur

The output of SAX algorithm can be interpreted as a series of symbols. Sequitur is an algorithm for inducing a hierarchical grammar from a given sequence of symbols, thereby compressing the input to a smaller representation [15]. This outcome small representation is, again, a series of symbols. Some symbols are *terminal*. A terminal symbol is a symbol that was also present in the original sequence. Other symbols are *non-terminal*. A non-terminal symbol is an outcome of the compression, and there is an associated *rule* which determines how to ‘expand’ this symbol to again a series of (non)-terminal symbols. A grammar is defined as a set of rules for repeatedly expanding all non-terminal symbols in this way will eventually reproduce the original sequence, with no more non-terminal symbols present. A special ‘start symbol’ (denoted by *S* in this text) determines the initial rule to use when starting the expansion. The Sequitur algorithm constructs a grammar from the input sequence by processing input symbols sequentially, and adding them to the grammar one by one. An example of an output grammar is given in Table 1. The symbol-to-rule mapping is given by *R* : symbol → sequence.

**Table 1:**
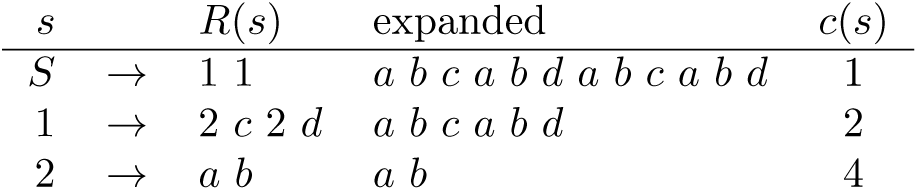
Example grammar produced by Sequitur for the input sequence *a b c a b d a b c a b d*, demonstrating rules, expanded sequences and occurrence counts. Note that *a, b, c, d* are terminal symbols, and *S*, 1 and 2 are non-terminal symbols in this example.

| $s$ | | $R(s)$ | expanded | $c(s)$ |
| --- | --- | --- | --- | --- |
| $S$ | → | $1\ 1$ | $a\ b\ c\ a\ b\ d\ a\ b\ c\ a\ b\ d$ | 1 |
| 1 | → | $2\ c\ 2\ d$ | $a\ b\ c\ a\ b\ d$ | 2 |
| 2 | → | $a\ b$ | $a\ b$ | 4 |

The Sequitur algorithm processes input symbols sequentially, adding them to the grammar one by one, while maintaining two grammar properties:

**digram uniqueness** : every pair of symbols (terminal or non-terminal) occurs *not more than once* in the grammar. If the same pair of symbols occurs twice at some point, a new rule is created mapping a non-terminal symbol to this sequence of two symbols, and the two occurrences of the pair are replaced by the new non-terminal symbol. This property ensures that the grammar gets compressed once a repetition is found. It also imposes a hierarchical structure on the grammar.
**rule utility** : every rule is used *at least twice*. Because of the digram uniqueness property, it may happen that a pair consisting of a non-terminal symbol and of another symbol is condensed as a new rule. This causes that non-terminal symbol’s occurrence count drops by one, which might become the only occurrence of the non-terminal symbol. In this case, the rule is removed, and its non-terminal symbol is replaced by the corresponding sequence. This property not only ensures that no useless rules (with only one occurrence) exist, but also allows for formation of rules longer than two symbols, which would otherwise not be possible.

By maintaining these properties, and using the proper data structures, the Sequitur algorithm runs in linear time in the length of the input sequence [15]. The output grammar can then be used to determine the relative frequency of sub-sequences by their occurrence count, so that they may be used in the quantification method.

In the context of finding common patterns in the SAX output, we are interested in the *occurrence count* of the different non-terminal symbols, since these represent common sequences. Given a grammar, let *s* denote a non-terminal symbol in that grammar. Then we can find the occurrence count *c*(*s*) of that symbol by finding all the rules that *s* occurs in, and taking the total occurrence count of the symbols associated with all those rules. *S* always has an occurrence of 1 (Equation 3).

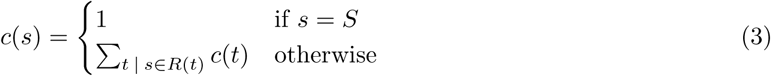

For example, in Table 1, the most frequent sub-sequence (longer than 1 symbol) is *a b*, as it occurs four times in the input. We introduced the occurrence count and added to the algorithm in order to apply Sequitor to this particular research problem, namely finding patterns in the SAX output data.

#### 2.2.3. Feature vectors

The *k*-motifs method, using SAX and Sequitur, yields frequent motifs for a certain time series, which may be used to give insights into rodent behaviour. However, these motifs are not suited for classification since they do not take the form of a feature vector. Specifically, there is no obvious comparison between motifs derived from different time series.

To resolve this issue, the following method is applied: given a set of time series from which feature vectors need to be extracted, first find the *k most interesting* motifs (defined hereafter) for the complete dataset. We further refer to this set of interesting motifs as ℳ_*k*_(*D*) (Equation 4). Now, in order to create a feature vector for a single time series, ℱ_*k*_(*d* ∈ *D*), we need to determine the *frequency* of each motif discovered in that time series (Equation 5). This method yields feature vectors that are comparable for classification, element-wise.

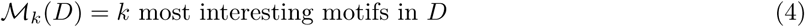

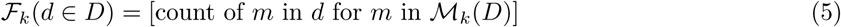

In the context of this manuscript, the classification relates to discrimination between the experimental intervention group compared to the control (vehicle treated) group (on the basis of open field data). These groups, in machine learning terminology, are referred to as *classes*. In many classification problems, the uneven size of classes is an issue. To account for this, *k* frequent motifs should be extracted from the data points of each class separately. Given *n* classes, this will yield *k* · *n* frequent motifs. As before, a feature vector representing *d* ∈ *D* is created by determining the frequency of each of the motifs in the data point. Thus, the length of the feature vector, is dependent on the number of classes *n* as well as the number of extracted motifs *k*.

#### 2.2.4. Interestingness

By means of combinatorics, the number of possible motifs grows very fast with the length of the sequence. Therefore, one should define some rules for motif preference. This can be achieved by defining ‘interestingness’ of motifs. Given a motif *m*, three interesting properties may be defined, each of which one may wish to maximise when searching for interesting motifs. To evaluate a motif’s interestingness *I*(*m*), some (non-linear) combination of these three properties can be used. The *k* most interesting motifs could then be selected as those with the highest interestingness.

**frequency** *I*_*f*_ (*m*) : the number of times this motif occurs in the data; the more frequent a motif is, the more suitable it will be as a feature. Of course, certain motifs which happen to be rare, can still carry meaningful information and be useful for classification but in practice, there is no efficient way to select these motifs because using rare motifs would facilitate over-fitting (i.e., building a classification tool that contains more parameters than can be justified by the data, and will fail to predict future observations).
**length** *I*_*l*_(*m*) = |*m*| : the number of symbols in the motif. Longer motifs are more interesting because they expose more detailed patterns. A motif of length 2 only shows a linear movement from one position to another, while longer movements will describe more detailed aspects of the animal’s behaviour.
**diversity** *I*_*d*_(*m*) : the number of unique symbols in the motif. A long motif may only contain few unique symbols, which means the motif describes a movement with a high amount of repetition. Since such a pattern is clearly a combination of other motifs, it might be more interesting to focus on motifs which cannot be easily decomposed. This is encouraged by preferring motifs with a high number of unique symbols.

Certain relations hold between these properties. Given any motif *m* and symbol *s*, a new motif may be constructed as *n* = *m* + *s* by appending *s* to *m*. Regardless of the choice of *m* and *s*, it is clear that *n* cannot be more frequent than *m*, since for each occurrence of *n* there is also an occurrence of *m* contained within. Thus, *I*_*f*_ (*m* + *s*) ≤ *I*_*f*_ (*m*) and |*m* + *s*| > |*m*| for any *m* and *s*. This means that frequency is inversely proportional to length: *I*_*f*_ (*m*) ∝ |*m*| ^− 1^. Note also that |*m*| ≥ *I*_*d*_(*m*), meaning that length must grow with diversity, and so *I*_*f*_ (*m*) ∝ *I*_*d*_(*m*)^−1^.

This means that, when using these three properties and *k*-motifs to define a quantification method, high accuracy (*I*_*f*_) and high interpretability (*I*_*l*_ and *I*_*d*_) are mutually exclusive. It is not clear whether this holds for quantification methods in general. In any case, the best way to combine the three properties into a single measure of interestingness *I*(*m*) is not obvious. In this work, four different ways of combining the three measures are explored (Equations 6-9). While *I*_1_ combines the three measures with equal weight, the other functions favour one measure while devaluing the other measures by scaling them logarithmically.

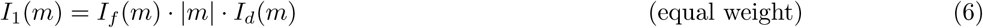

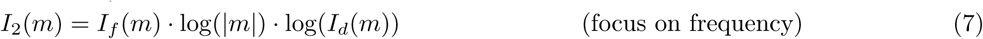

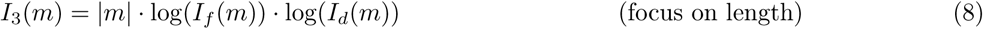

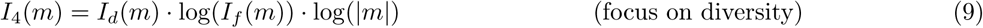

The choice of combination matters because they can result in different sets of leading motifs (Fig. 5).

#### 2.2.5. Spatial relations

Although the *k*-motifs algorithm may be applied to any time series, only the *X centre* and *Y centre* variables from *V* are used to construct the feature vectors. The choice to restrict the number of variables was made in order to reduce the size of the feature vector, thereby increasing interpretability. The choice for these specific variables was made since they are the variables used by other methods, e.g., T-patterns. Then, we should account not only for the subject’s position in space, but also for the relationship to *objects* and to the *boundaries* of the open field. Therefore, the *k*-motifs algorithm must not only be applied to a time series tracking the subject’s position, but also to time series reparameterised in relation to objects. Each of these transformed time series is called a *spatial relation* (including the original position).

This extension of the feature space further increases the length of the feature vector to *k* · *n* · *r*, with *r* the number of relations we wish to capture. The spatial relations that we chose, are presented in Table 2 (although of course, more possibilities exist). In our specific dataset, *n* = 2 (control versus OCD) and *r* = 4 (absolute position, relative position, minimum distance to any object, minimum distance to the boundary). The length of the feature vector, then, becomes 8 · *k*.

**Table 2:** Chosen spatial relations.

| Name | Dimensions | Description |
| --- | --- | --- |
| absolute position | 2 | $(x, y)$ location of the animal on the table in Cartesian coordinates |
| relative position | 2 | change in $(x, y)$ location of the animal as compared to the previous time point |
| nearest object | 1 | the distance from the animal to the nearest object |
| nearest boundary | 1 | the distance from the animal to the nearest point on the boundary of the table |

Note that some spatial relations are two-dimensional (absolute / relative position) while others are one-dimensional (distance). While using the *k*-motifs algorithm in application to one-dimensional data is straightforward, in case of two-dimensional data it becomes is less obvious. We took the following approach. First, the two dimensions are processed by SAX individually, resulting in two separate symbolic time series. These two time series are then recombined to create a time series consisting of pairs of symbols. This new time series can then be processed by Sequitur, interpreting each pair as an individual symbol.

### 2.3. Free parameters

The complete quantification method, combining *k*-motifs, creation of the feature vector and spatial relations, involves a number of free parameters (Table 3).

**Table 3:**
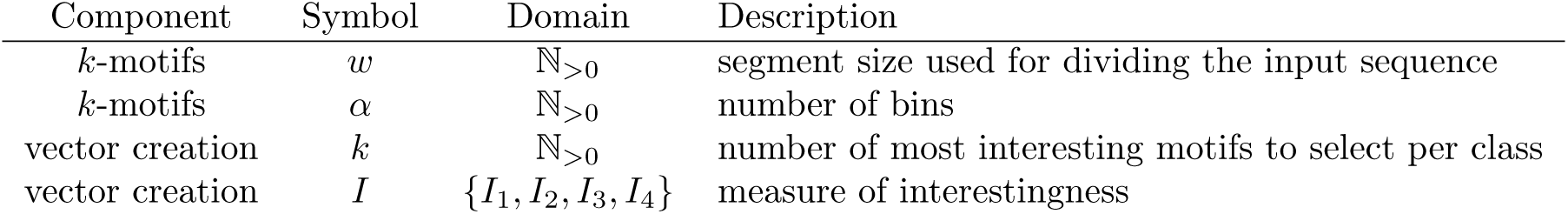
An overview of the free parameters for the full *k*-motifs pipeline.

Hill-climbing is a technique for optimising a cost-function characterising certain problem, by iteratively adjusting its input parameters [16]. This technique can be used to determine the optimal values in the parameter space (*w, α* and *k*), whereas exhaustive search is performed to optimise *I*. In order to evaluate the results of this optimisation, the complete dataset is split into two portions: 90% that is used for hill-climbing based on classifier performance (which in turn splits the data into training and test sets) and 10% that is held out for final evaluation.

To summarize, the schematic representation for the full *k*-motifs pipeline is given in Figure 3.

**Figure 3:**
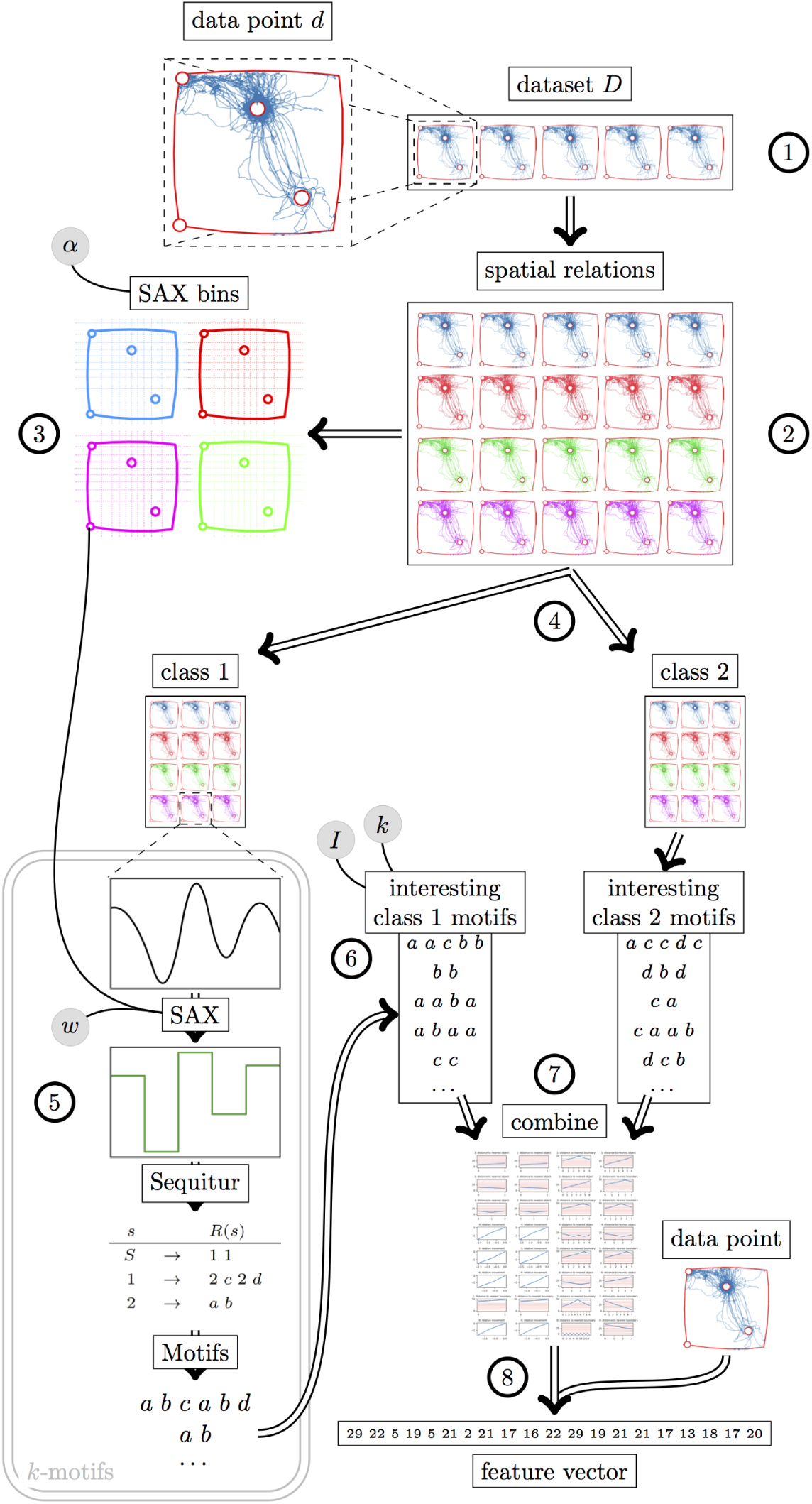
Schematic visualisation of the approach proposed in this work, based on *k*-motifs. The pipeline starts with the raw dataset (1), containing movement data from multiple open field sessions. Each of these sessions is then transformed into multiple spatial relations (2). For each spatial relation, the SAX bins are computed, using parameter *α* determining a total number of bins (3). The spatial relations are then separated based on the class of the session (4). For each class, every data point is processed using *k*-motifs, using the parameter *w* and the SAX bins computed earlier (5). For each data point, this results in a set of motifs, and all motifs of a class are gathered into a list. The motifs are sorted according to a measure of interestingness (*I*). Subsequently, certain number *k* of top motifs is selected to represent the class (6). The top motifs from each class are then combined to create the final set of motifs (7). These can then be used to create a feature vector for any data point (from the original data or otherwise), by counting the number of occurrences of each motif within that segment (8).

### 2.4. Evaluation

Four popular classifiers are used to compare four different quantification methods. Implementation of these classifiers comes from sklearn package for Python [17]:

**Gaussian Naive Bayes** Assuming independence between features, Bayes’ theorem can be applied to the data. This method learns probabilities with each combination of class and feature from the data, and from this evaluation, the most probable class can be inferred for a new observation[18].
**Decision Tree** In this method, a number of if-then-else decision rules are learned with the purpose to accurately divide the data into classes based on feature values. A new observation is then classified by evaluating the decision rules on its features [19].
**Multilayer Perceptron** A feed-forward network of artificial neurons with nonlinear activation functions is used to model the relationship in the data between features and classes through connectivity weights between neurons. A new observation is then classified by feeding a set of features representing this observation into the network, which returns the predicted class at the output [20].
***k*-Nearest Neighbours** All observations in the data and their associated classes are stored in memory. Given a new observation, its class it determined by majority vote of the *k* data points closest to the observation in the feature space [21].

We chose the default parameters as implemented in the sklearn package. In order to determine the relative performance of classifiers, the label-frequency based macro *F*_1_ score was used as a measure of test accuracy. As a variation of the commonly used *F*_1_ score, this measure combines the classifier’s precision *P* (also known as positive predictive value, PPV) and recall *R* (also known as sensitivity) with equal weights in order to evaluate the performance of the classifier. The *F*_1_ score is computed individually for each class *c*, then the average between these scores is computed, with each score weighted by the size of the corresponding class |*c*| (Equation 10) [22]. This ensures that imbalance in class size does not influence the score.

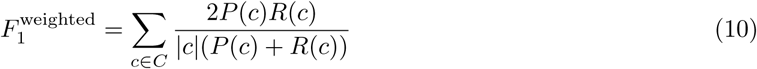

In order to obtain reliable classification results, stratified *k*-fold cross-validation was used. In non-stratified *k*-fold cross-validation, the data is randomly divided into *k* subsets of equal size. Then, for each subsample *i*, the subsample is used as a test set while the other *k* − 1 subsamples are used for training. This results in *k* classifiers with each a classification score, which may be averaged to obtain the final score. This method ensures that all individual data samples are used for testing exactly once. Additionally, in *stratified k*-fold cross-validation, each subset is selected in a way that the distribution of classes is as near as possible to that of the complete data [23] (here, it means a ratio 1:1 between cases and controls). This prevents confounding the results by the class labels. In this project, *k* = 10 is used.

### 2.5. Comparison with other methods

In this work, we further compare k-motif approach with other methods which also define how a single experimental session is reduced to a simpler representation:

1. established T-pattern method [8] which interprets patterns as sequences of visited discrete fields (in this case, on the square experimental table divided into 25 square zones) within certain time-frame
2. *full data* method: no reduction is performed and the complete set of points visited during the session is taken as the representation of that session
3. *means and variances* method: the data of a session is reduced to just four numbers: the mean and the variance, per dimension (one of each for x-coordinates and y-coordinates)
4. *behavioural* method: the data of a session is reduced by applying several hand-crafted heuristics to it that are thought to be useful descriptions of an animal’s behaviour. This results in a collection of numbers that describe behaviour displayed during the session. For a full description of this method, check [24]

sectionResults

### 2.6. Normalising bins with SAX algorithm

Using the T-pattern method with linear zone divisions, the entropy of the proportion of visits per zone is 2.90. In contrast, using the bin divisions based on SAX algorithm, the entropy is 2.63 (Figure 4 A).

**Figure 4:**
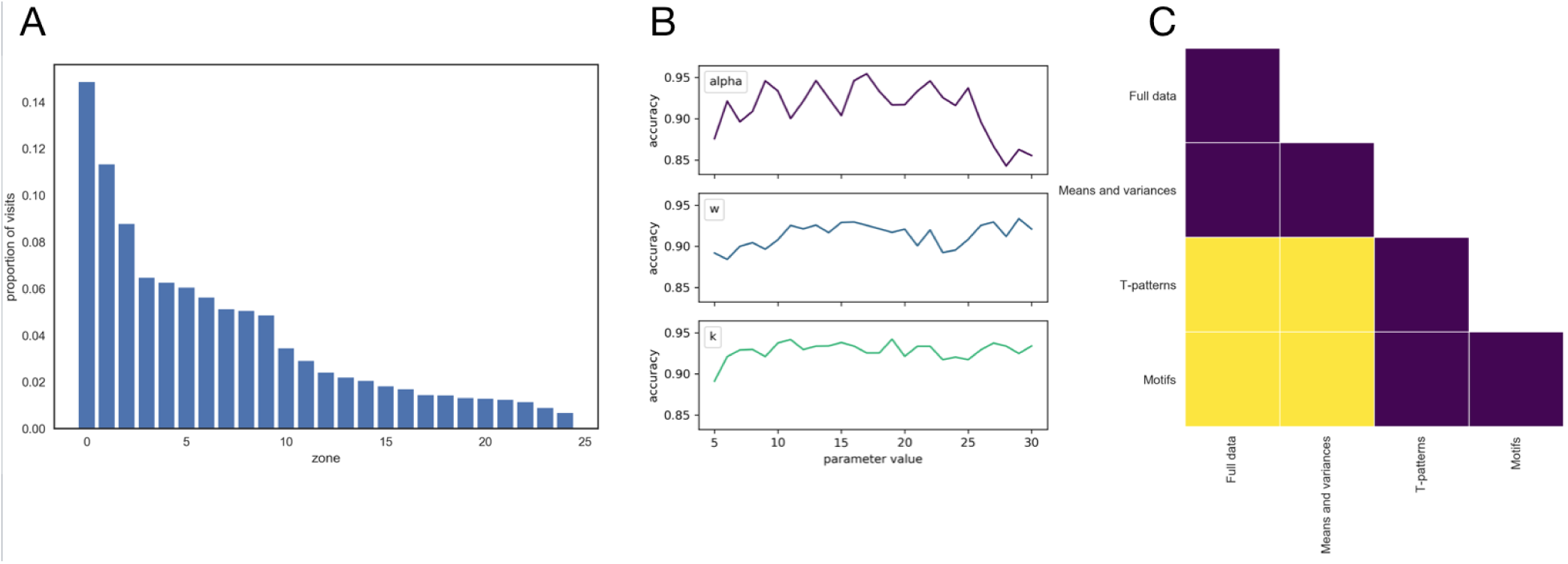
**A**: Proportion of visits to each bin in the ‘2014’ dataset after using SAX with parameters *α* = 5 and *w* = 15. **B**: Robustness of the k-motifs algorithm using k Nearest Neighbors using *I*_2_ definition of interestingness, with respect to parameters *α, w* and *k*. **C**: Significance of differences between method performances. Yellow fields denote significant differences in performance between two corresponding methods, while purple means non-significant.

### 2.7. Leading motifs depending on interestingness measure

The 8 most interesting motifs obtained with use of every of the four measures of interestingness defined in Section 2.2.4, are presented in Figure 5. An example of a feature vector based on these most interesting motifs is presented in Table 4.

**Figure 5:**
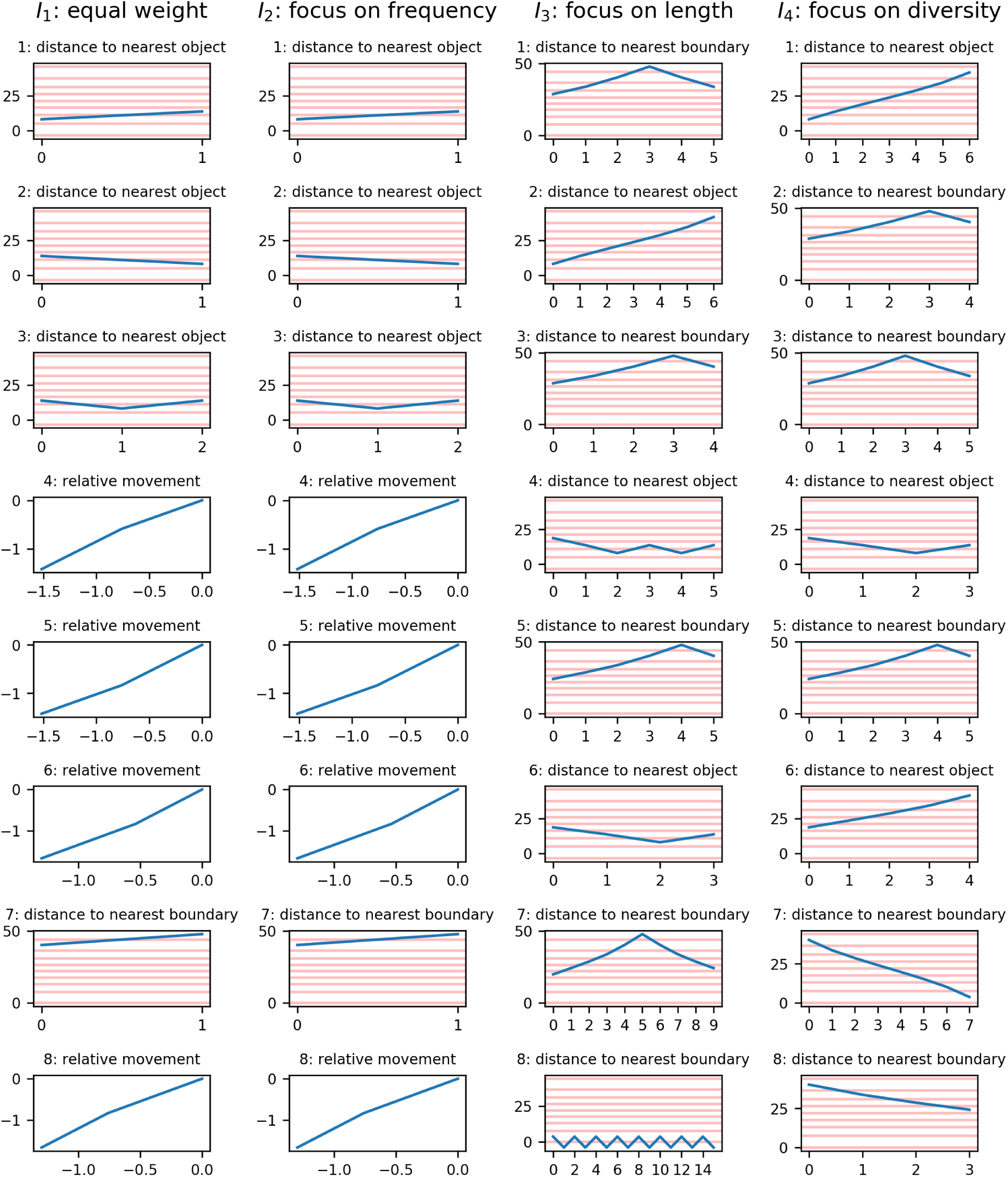
8 most interesting motifs resulting from the four different combinations of the three measures of interestingness as defined in Section 2.2.4. Note that one-dimensional features (distances) are presented in time scale, while two-dimensional features (movement) are shown in space. Relative movement patterns originate at the central position (0, 0). Where possible, bins (i.e. zones) are indicated with red lines (this is not possible for relative movement as the bins re-orient with each step).

**Table 4:**
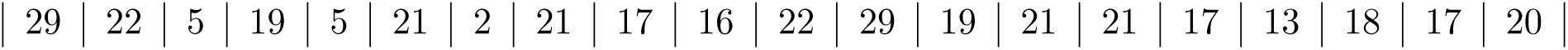
An example feature vector resulting from the quantification method, describing a single data segment. This vector contains 20 features, since there are two classes and *k* = 10, so 10 motifs from each class are used. Each number indicates the number of times a particular motif occurs in the data segment of interest.

A summary of all 20 (2 classes × 10 motifs) feature vectors obtained using the *I*_2_ measure for all the data segments in our dataset (concatanated for the whole cohort) can be found in Figure 6. The hill-climbing method found *w* = 15, *α* = 10, *k* = 10 and *I* = *I*_2_ as parameters yielding the highest classification performance. Note that *k* = 10 was the highest allowed value in the hill-climbing setup. This suggests that segments of 625 [ms] in length and 10 bins per dimension should be used as an input to the SAX algorithm. Further, top 10 motifs should be delegated to represent each class and since *I*_2_ was the leading method for quantifying interestingness, motifs should be chosen primarily due for their frequency, rather than due to their length or diversity. Also, performance accuracy is robust with respect to *α, w* and *k* (Figure 4 B).

**Figure 6:**
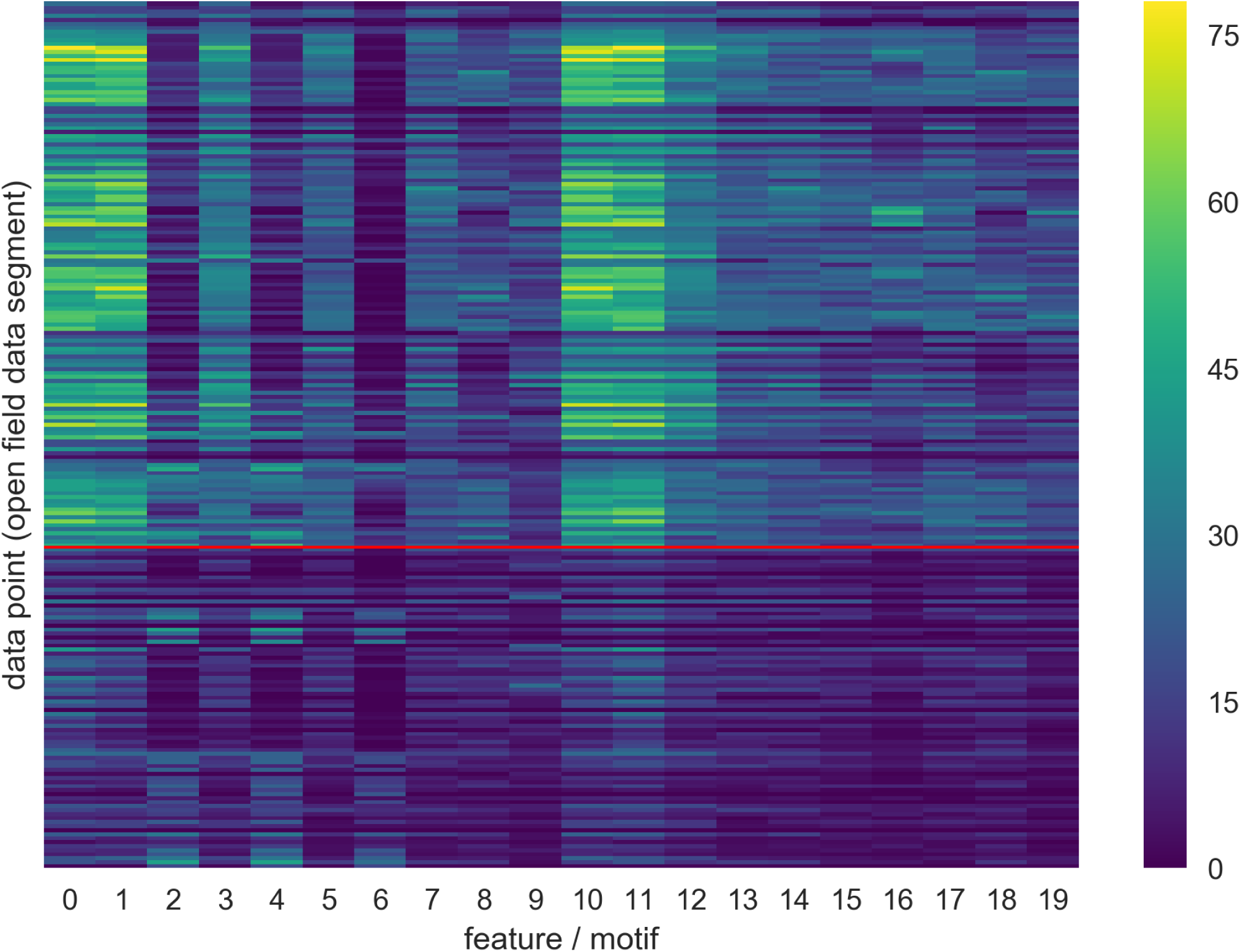
A summary of all feature vectors for all data points in the *2014* dataset, using the *I*_2_ measure. Note that data points are separated by class, as indicated by the red line (with the quinpirole group at the top and the control group at the bottom).

### 2.8. Comparison with other methods

The full summary of the performance of different classifiers can be found in Table 5. The new quantification method was able to achieve an average score of 0.86, and as such, it outperforms all of the other classifiers. Note that although the average classification score is the highest, not all the individual scores are the highest. In particular, the MultiLayer Perceptron classifier did not yield a high score for the k-motif method (0.69), while T-patterns scored high in that case (0.84). The highest score of any method-classifier combination was achieved by the k-motif method using *k*-Nearest Neighbors (0.94).

**Table 5:**
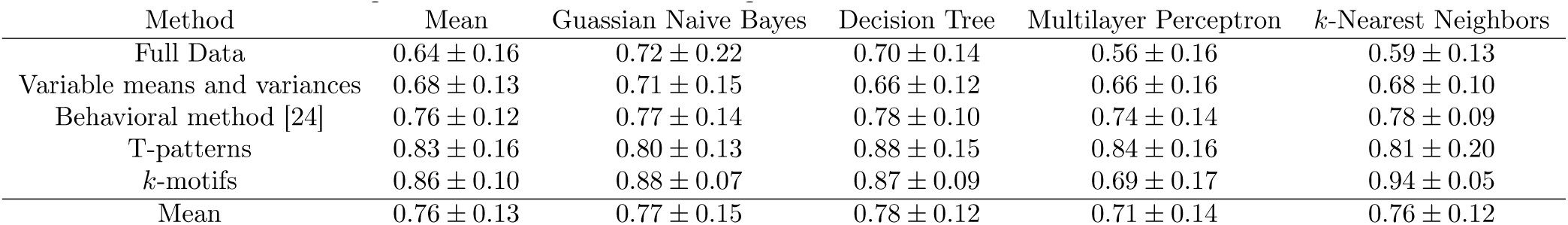
Classification performances of the different quantification methods for each of the considered classifiers.

In this classification study, the validation was performed using stratified 10-fold cross validation (implemented with use of *sklearn* Python package).According to our results, the k-motif approach is significantly better than using full datasets, and using means and variances from raw experimental variables, but not significantly better than either the behavioural method [24] or the T-pattern method [8]. Significant differences between methods are visually presented in Figure 4 C.

## 3. Discussion

The current approach examined the utility of the k-motif approach as a means to improve automated behavioural analysis in the open field. Our results suggest that this is an efficient technique which returns a sparse set of behavioral patterns best discriminating between experimental groups and gives higher classification performance than the available methods.

### 3.1. Normalising bins

The results were contrary to expectations, meaning that dividing the table into linear zones gives better performance than normalising divisions in creating a uniform distribution of zone visits. However, it is unclear whether this effect is specific to this particular dataset and additional validation with other datasets would be useful. While the literature suggests that normalising bins should aid in creating uniform visit distributions, it would be interesting to investigate the method further in other datasets.

### 3.2. Motifs, feature vectors and interestingness

The motifs yielded by *k*-motifs were interesting, although they may be difficult to interpret in a systematic way. Most probably, motifs are specific to the particular shape and orientation of the open field arena used and the associated objects, and in another experimental setup, another set of motifs might come out as most predictive of the subject class. The k-motif approach, in the form proposed in this work, is an exploratory technique but is adaptable to different contexts. One noticeable property of motifs obtained in this study is that most of them, describe mostly linear motion or repetitions of linear motions. For example, many of the motifs in Figure 5 show a single oscillation of a repeating motion, or else they show an almost linear movement. These movements are not very complex, and may, perhaps, be also picked up by a more constrained and computationally cheaper method. Some of the features of the motifs presented in Figure 5 are particularly interesting. Of note, the motifs obtained from interestingness *I*_1_ and *I*_2_ are identical. Since the difference between these measures lies in the way that length and diversity of motifs are valued, while the value put on motif frequency is constant, this suggests that even in measure I (the equal-weight measure) the frequency has more influence on the final selection. It may be useful to normalise the three properties before combining them into one measure, so that any of them has equal range of values, before taking the sum.

Furthermore, the nature of each measure of interestingness is reflected in the presented motifs. Both the measure focusing on length and the measure focusing on diversity do produce longer motifs, while the measure focusing on diversity additionally finds motifs with a larger number of unique values. Interestingly, multiple motifs are shared between different measures of interestingness focusing on length *I*_3_ and diversity *I*_4_, while seemingly no motifs are shared between these and the measure focusing on frequency *I*_2_. This can be explained by the fact that frequency of motifs should be inversely proportional to both motif length and diversity. It should be noted that the most-interesting motifs presented in Figure 5 include motifs of all relation types (positions and distances) except for the ‘relative position’ type. This means that no motif of this type was among the 8 most interesting based on any of the four interestingness measures. This may be explained by the fact that the relative position may have the highest variance of all of the spatial relations, meaning that patterns are less likely to occur.

From the classifier performance results, it would appear that *I*_2_, the measure focusing on frequency, is the best for determining the interestingness of a motif. Note that the significance of the differences between measures was not evaluated. The goal of exploring different measures was not only to increase performance, but also interpretability, and this is one of the reasons we proposed the criteria *I*_3_ and *I*_4_. However, motifs produced by *I*_3_ and *I*_4_ seem at most marginally more interpretable than the others. Our conclusion is that, in this case, it would be best to focus on *I*_2_, as properties other than motif frequency do not yield any additional benefits to the ability to classify behavioural patterns.

Figure 6 demonstrates some interesting properties of the feature vectors. The data points are separated by class along the vertical axis, and a clear separation is visible in the values of the feature vectors. This visually shows the source of the classification accuracy as found in the evaluation. However, the main difference between the two groups seems to be that many motifs are frequent in the quinpirole treated group, while few motifs are frequent in the control group. Although this facilitates classification, it is contrary to expectations: since 10 motifs were taken from the top motifs in each class, the expectation would be for half of the motifs to be frequent in all data points of one class, and the other half in the other class. One possible explanation for this phenomenon is the fact that motifs derived from the control class, although common in the control class, are even more frequent in the quinpirole treated class.

### 3.3. Parameters

Parameters optimized in the hill climbing algorithm specified twice as high number of bins per dimension, than it is specified in the T-pattern method. It would be interesting to explore how the T-pattern method would perform if using a higher number of bins. Although *k* = 10 was the optimum value, this was also the highest allowed value for *k* in the chosen parameter range. If higher values were allowed, it is likely that the performance would keep increasing with *k*, since an increase in the number of features can only increase the classification performance. However, a higher value of *k* would also negatively impact the computational complexity of the algorithm. In addition to the four aspects of the method that were parameterised, some more possibilities exist that were not explored, e.g., a choice of which spatial relations to include. Also, interestingness of a motif could be defined in multiple other ways.

### 3.4. Evaluation

Our results indicate that - although the *k*-motif method performs significantly better than methods based on full data, and data compressed to means and variances - it does not perform significantly other than either of the two other, more complex methods. However, it should be taken into consideration that the parameters of the k-motif method were fine-tuned to our dataset, while the other methods were not privileged in a similar fashion. This manifests in the fact that the performance on the held-out data was only 0.75. The performance was lowest for the Multilayer Perceptron classifier, which is known to be sensitive to overfitting [25, 26], which could also contribute to this effect. In the future, cross-validation of the results on another experimental cohort might be performed, although each cohort behaves differently in the open field environment and the classification performance would certainly drop.

One aspect of classification that was not explored in this project, but which would be very interesting for the future, is the evaluation of the relative contributions of features in the process of classification. It would be interesting to know which motifs, or which spatial relations, contribute most to determining classes in order to refine the k-motif algorithm further.

### 3.5. Conclusion

The new quantification method based on *k*-motifs, which converts patterns discovered by k-motifs into feature vectors, performs very well on the open field datasets arising from the repeated checking behaviour resulting from quinpirole administration in rats. This allows for an additional description of behaviors characteristic to the experimental group, as opposed to the control group.As such, it can serve as an alternative to existing methods dedicated to quantification of animal behaviour in the open field such as the T-patterns algorithm.

## Author Contributions

MK, NB, KK and JG designed the study. KK performed the open field experiments and was consulting the team with respect to standard methods for quantifying behavior in the open field. MK and NB conducted the computational study and drafted the manuscript. NB, MB and JG co-supervised the MK’s Bachelor Thesis project. JG, MB, TC and JB critically revised the draft of the manuscript.

## Acknowledgments

We would like to thank the members of the **Translational Psychiatry group** (PI: Jeffrey Glennon) for sharing their expertise on behavioral paradigms in rodents, and to the Department of Artifical Intelligence, Radboud University Nijmegen, for facilitating this research as a topic for MK. We would also like to thank **Jesse Zwamborn**, within the Radboud University AI department for his contribution to this project at the early stages.

## Ethical approval

The animal experiments in rats with use of quinpirole, were carried out in accordance with local guidelines and regulations within the Netherlands. All animal procedures were approved by the Ethical Committee on Animal Experimentation of Radboud University Nijmegen (under RU-DEC, number 2012-281) and the Dutch Ethical Committee on Animal Experimentation, and in accordance with the EU guidelines for animal experimentation.

## Data availability statement

All the codes developed for this manuscript, are publicly available at https://github.com/MareinK/kmotifs-paper-code.

The behavioural datasets are handled by the TACTICS consortium and are not publicly available, and not published. For access to this data, please contact Dr Jeffrey Glennon at.

## 4. Appendices

### 4.1. Experimental datasets

**Figure 7:**
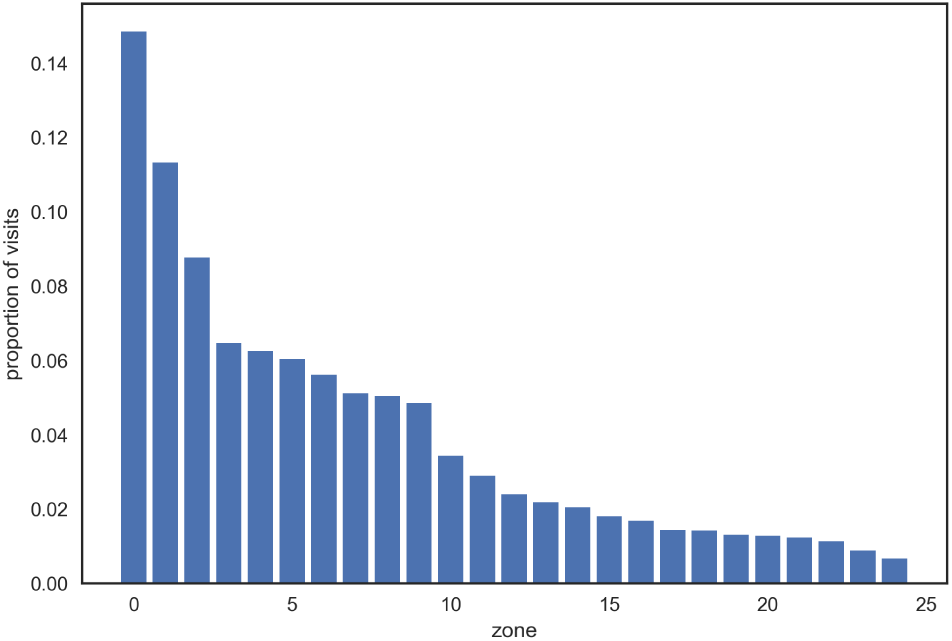
Proportion of visits to each zone in the open field, in our datasets (Section 2.1). The distribution is significantly non-uniform (Kolmogorov-Smirnov test at *p* < 0.05), which breaks the assumptions of the T-pattern method.

**Table 6:** Descriptions of experimental variables recorded with Ethovision software.

| Variable | Unit | Description |
| --- | --- | --- |
| Trial time | seconds | time since the animal was introduced into the open field environment |
| Recording time | seconds | time since the video recording was started |
| X centre | centimetres | x-coordinate of the animal's centre |
| Y centre | centimetres | y-coordinate of the animal's centre |
| X nose | centimetres | x-coordinate of the animal's nose |
| Y nose | centimetres | y-coordinate of the animal's nose |
| X tail | centimetres | x-coordinate of the animal's tail |
| Y tail | centimetres | y-coordinate of the animal's tail |
| Area | square centimetres | the amount of area covered by the body of the animal |
| Area change | square centimetres | the amount of body area that does not overlap with the body area of the previous measurement [27] |
| Elongation |  | a measure of how much the animal stretches its body out, between 0 and 1 |
| Direction | degrees | the direction the animal is facing |

### 4.2. Data preparation

The datasets were incomplete, which is often the case in translational psychiatry experiments. It was found that the experimental data contained missing values, non-uniform structure, redundancy and anomalous data. For each of these issues, a solution was proposed and implemented. Most solutions involve disregarding parts of the original data, but enough data remains to support the aim of the project, especially after augmentation (Section 4.3).

#### 4.2.1. Missing values

Some sessions were missing while some other sessions were cut short, as a result of technical problems that occurred during the experiment. Lastly, at certain time points within sessions, some data was missing, leading to long contiguous periods missing data within a session. Specifically, there were some gaps of a spatial nature, occurring in multiple sessions, always at the same area in space (Figure 8). In sessions affected by these gaps, whenever a subject passes through the affected area, a temporal gap in the data occurs. Since the spatial form of these gaps is roughly elliptical, these kinds of gaps are most likely caused by shadows or highlights on the surface of the open field. These features are caused by the computer vision software determining the subject’s position. The effect is less prominent in the *X centre* and *Y centre* variables, with most missing values occurring in the other *X* and *Y* variables.

**Figure 8:**
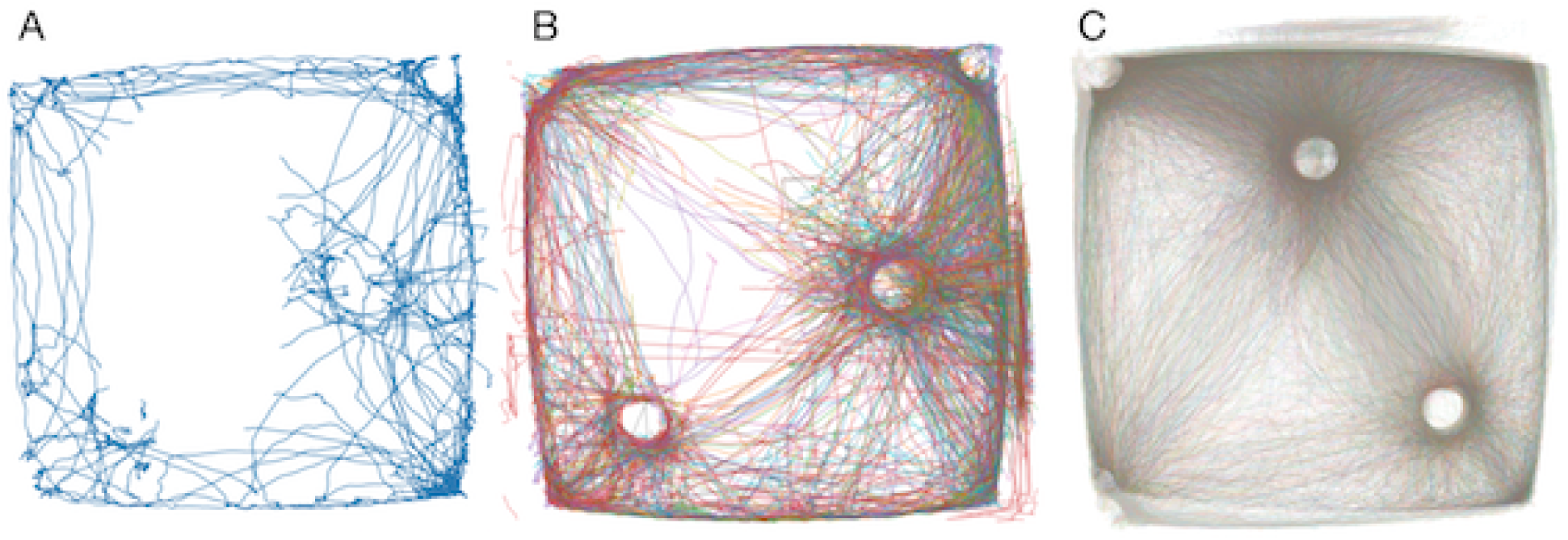
**A**: An exemplary case of missing values of a spatial nature, a single session (this data was not used in the final study). A large area to the centre left of the open field is devoid of paths; all encroaching paths are cut off at the boundary of the area. **B**: The issue of missing values of a spatial nature, an overlay of a few sessions, with each subject marked with a different colour (this data was not used in the final study). The gap, although less pronounced, is also visible here, showing that it is a structural problem. **C**: An overlay of all movement data for all sessions from two different datasets: the dataset used in our study, and another dataset produced using the same experimental setup. Although the locations of objects and boundaries are aligned between sessions within datasets, there is a clear difference in these locations between datasets, as can be seen by the displacement effect in the image. The projections from the two datasets have been translated and rotated to achieve a best fit, but even so the difference is clear.

Multiple solutions to these problems were considered, most prominently either interpolating the data to fill the gaps, or complete removal of sessions containing missing data. Interpolation was difficult in this case, as this would require prior characterisation of the existing data, while characterisation was the point of the method. Hence, we decided to remove all the sessions containing missing data (Table 7).

**Table 7:** Sessions retained after pruning the dataset. The group to which each rat belongs (*s*aline or *q*uinpirole) is presented in the second row. The leftmost column presents the session number. The inner cells show the internal identifier for each rat-session combination.

| rat | 109 | 110 | 111 | 112 | 113 | 114 | 115 | 117 |
| --- | --- | --- | --- | --- | --- | --- | --- | --- |
| grp. | s | s | s | q | q | q | q | q |
| s1 | 4 | 5 | 6 | 7 | 8 | 9 | 10 | 12 |
| s2 | 16 | 17 | 18 | 19 | 20 | 21 | 22 | 24 |
| s3 | 28 | 29 | 30 | 31 | 32 | 33 | 34 | 36 |
| s4 | 40 | 41 | 42 | 43 | 44 | 45 | 46 | 48 |
| s5 | 52 | 53 | 54 | 55 | 56 | 57 | 58 | 60 |
| s6 | 64 | 65 | 66 | 67 | 68 | 69 | 70 | 72 |
| s7 | 76 | 77 | 78 | 79 | 80 | 81 | 82 | 84 |
| s8 | 88 | 89 | 90 | 91 | 92 | 93 | 94 | 96 |
| s9 | 100 | 101 | 102 | 103 | 104 | 105 | 106 | 108 |
| s10 | 112 | 113 | 114 | 115 | 116 | 117 | 118 | 120 |

#### 4.2.2. Non-uniformity

In this work, we had a selection of datasets to work on, coming from a few distinct experiments on the quinpirole model of obsessive-compulsive disorder in rats. However, the data displayed problematic non-uniformity between and within datasets, causing problems when comparing data points. For example, the scale and orientation of the coordinate space might differ between datasets. These problems can largely be attributed to differences in camera position and angle at the time of recording. Because of perspective and lens distortion effects, these camera parameters have a large effect on the shape of the open field as projected into two-dimensional space. Although within any single session the camera is stationary, these effects cause problems when attempting to compare between sessions and datasets where the camera parameters differ (Figure 8 C). For this reason, only a single dataset is used for the project, namely the dataset which contains the largest number of sessions, and also happens to display uniform camera parameters between sessions, was used for the final study.

An additional problem is caused by the fisheye effect (Figure 8). An almost unpreventable effect of using a video camera, it causes the recorded footage to become warped. Since Ethovision 3.0 does not take this issue into account, the resulting data is also warped. Although the severity of the effect on further data analysis is not obvious, it would be good to try and revert the warping effect in the resulting data. However, that has not been attempted in this project.

#### 4.2.3. Anomalies

In some sessions, some anomalous sudden jumps seemingly made by the subject over large distances from one location to another (the ‘teleportation effect’) can be found. Sometimes these anomalies even seem to transport the subject to locations outside the regular bounds of the open field (e.g., Figure 8 A, B). These jumps almost always come in pairs, where the subject is transported to an anomalous location and then transported back to the original area after a fraction of a second. It seems that these anomalies can be attributed to errors made by the computer vision software when extracting the movement data from the raw video footage. Anomalies may be detected by looking for position change between two adjacent time points that would be impossible for an actual rat to accomplish - assuming a top speed of 9.6 km/h for rats [28], 93 such event occur in the dataset (0.002% of all time points). One elegant solution to the problem of anomalies would be to detect these anomalies, remove the data between two instances of ‘teleportation’ so that the period of anomalous movement is no longer present, and then reconstruct the missing values by means of interpolation. However, as discussed in Section 4.2.1, interpolation is difficult and can cause that the classification problem becomes ill-posed. Therefore, since only such a small percentage of all data is affected with this problem, we decided to neglect it in this work.

#### 4.2.4. Redundancy

Since the original data contains a large number of variables, some of which describe phenomena that seem closely intertwined, there is redundancy in the data. For example, the location of the subject’s head can be predicted from the location of the subject’s torso with a high accuracy. It might then be possible to remove certain variables from the data, without losing predictive power in the classification. Therefore, we calculated pairwise correlations between all of the variables in the complete dataset (preprocessed with the previous step) with use or Pearson’s r (Figure 9 A).

**Figure 9:**
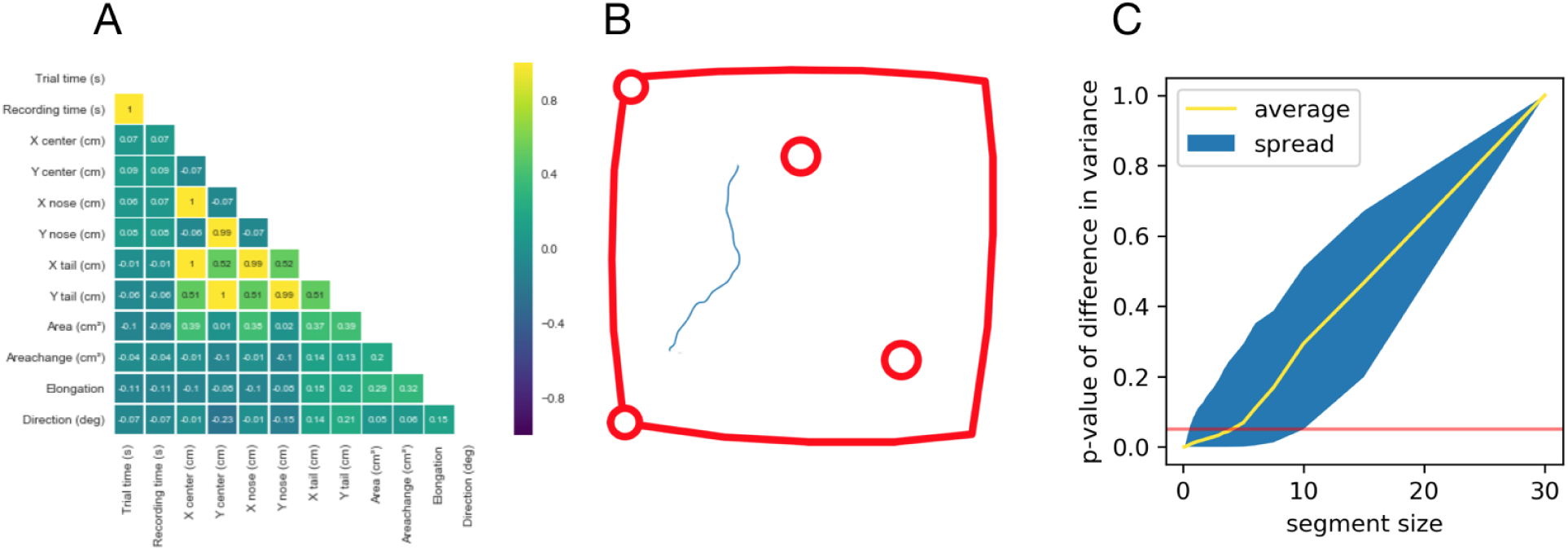
**A**: Correlations between variables recorded by Ethovision. **B**: a very small segment of an open field session) simply has too little features to use for describing behaviour. Compare with the images in Figure 1. **C**: Data segmentation. As the segment size decreases, so does the p-value of the the difference in variance as compared to the original segment size (30 minutes). The red line indicates a threshold *p* = 0.05. The yellow line presents the average p-value over all variables in *V*. The blue area shows the spread of the p-value for all variables.

Going further, the set of variables can be pruned so that only those variables remain which do not significantly correlate with any earlier variable. Given two variables *v, w* ∈ *V*, the variable *w* is ‘earlier’ than *v* (*w* < *v*) if *w* precedes *v* in the standard order of variables as used in Section 2. The resulting set of variables *V*′ is defined in Equation 11.

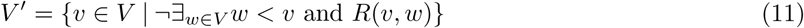

where *R*(*v, w*) - Pearson’s r coefficient between variables *w* and *v*. This pruning technique ensures that out of all sets of inter-correlating variables, only one remains, namely the first in the order as defined.

### 4.3. Data augmentation

Given the ultimate goal of creating a quantification method and evaluating it using classification, it is important to have sufficient data points for effectively training a classifier. Therefore, since the data undergoes significant reduction in the preprocessing stage, the possibility of data augmentation was considered. In machine learning, data augmentation is the process of extending the available data to be more multitudinal or vibrant, before using it as input for learning algorithms [29].

In this work, we augment the data. Given that each session is reduced to a single data point (set of features), the amount of data points is equal to the amount of sessions. However, each session has a duration of half an hour but the features are observables which be measured within far less time. Thus, each session is split into a number of segments, which can then all be used for feature extraction, yielding a larger number of data points.

Optimal split of the data into segments is not an trivial problem: if the sessions are split into too many segments, each segment will be so small for the proper feature estimation, while at the same time, the more splits, the more data points. Thus, there is a trade-off between the length of the data in each segment, and the number of segments (Figure 9 B, C).

Two methods for determining the optimal segment size are evaluated. The first method is based on the variance of the data in a segment as the segment size changes: as the segment size decreases, the variance in the data must decrease as well. The significance of this decrease from the original is calculated. Then, based on significance, the optimal segment size can be determined as a segment as small as possible while not displaying a significant difference in variance from the original data.

The second method for determining the optimal segment size is based on prior literature [4, 30, 31]. Many rodent experiments involve measuring the frequency of certain behaviours displayed by the animal, such as grooming or marble-burying. If the frequency of such behaviours can be determined from the literature, especially with regards to OCD behaviour, then this would be a good indication of how small segments can be while retaining the possibility to detect these behaviours in each segment.

## Footnotes

1 This may be evaluated by comparing the entropy of the visit counts, with a higher entropy meaning a more uniform distribution

